# Impulsivity and Emotional Dysregulation Predict Choice Behavior During a Competitive Multiplayer Game in Adolescents with Borderline Personality Disorder

**DOI:** 10.1101/2021.02.11.430783

**Authors:** A.C. Parr, O.G. Calancie, B. Coe, S. Khalid-Khan, D.P. Munoz

## Abstract

Impulsivity and emotional dysregulation are two core features of borderline personality disorder (BPD), and the neural mechanisms recruited during mixed-strategy interactions overlap with frontolimbic networks that have been implicated in BPD. We investigated strategic choice patterns during the classic two-player game, Matching Pennies, where the most efficient strategy is to choose each option randomly from trial-to-trial to avoid exploitation by one’s opponent. Twenty-seven female adolescents with BPD (mean age: 16 years) and twenty-seven age-matched female controls (mean age: 16 years) participated in an experiment that explored the relationship between strategic choice behavior and impulsivity in both groups and emotional dysregulation in BPD. Relative to controls, BPD participants showed fewer reinforcement learning biases, increased coefficient of variation in reaction times (CV), and more anticipatory decisions. A subset of BPD participants characterized by high levels of impulsivity and emotional dysregulation showed increased reward rate, increased entropy in choice patterns, decreased CV, and fewer anticipatory decisions relative to participants with lower indices, and emotion dysregulation mediated the relationship between impulsivity and CV in BPD. Finally, exploratory analyses revealed that increased vigilance to outcome was associated with higher reward rates, decreased variability in SRT, and fewer anticipatory decisions. In BPD, higher levels of emotion dysregulation corresponded to increased vigilance to outcome, and mediated its relationship with choice behavior. Together, our results suggest that impulsivity and emotional dysregulation contribute to variability in mixed-strategy decision-making in BPD, the latter of which may influence choice behavior by increasing attention to outcome information during the task.

## Introduction

Borderline Personality Disorder (BPD) is characterized by maladaptive decision-making tendencies, such as unstable relationships, self-harm behaviors, and substance use (for review, see Sebastian et al., 2014; Dougherty et al., 2004; Rosval et al., 2006; Soloff et al., 2000). Dimensional approaches propose that the symptoms of BPD reflect instantiations of an underlying predisposition toward impulsivity, emotion dysregulation, and interpersonal dysfunction (Allen & Hallquist, 2020; Beeney et al., 2018; Chapman et al., 2008; Hallquist et al., 2018; Hallquist & Pilkonis, 2012; Scott et al., 2014), which are transdiagnostic processes that contribute to maladaptive behaviors across several personality pathologies (see Allen & Hallquist, 2020 for a review). Personality and related psychiatric disorders, including BPD, often emerge and intensify during the adolescent period (Johnson, Cohen, & Brook, 2000; Johnson, Cohen, Smailes, et al., 2000; Larsen & Luna, 2018). However, due in part to extant controversy surrounding diagnoses of personality disorders in adolescence (Barker et al., 2015; Berenson et al., 2016; Chanen & McCutcheon, 2013; Coffey et al., 2011; Ka et al., 2008; Krause-Utz et al., 2016; Maraz et al., 2016) and the continued reliance on categorical classifications in the Diagnostic and Statistical Manual of Mental Disorders (DSM-5; American Psychiatric Association, 2013; Allen & Hallquist, 2020; Clark, 2007; Trull & Durrett, 2005; Widiger & Simonsen, 2005), little is known about how BPD symptoms emerge within the context of brain maturation, hampering the ability to develop effective preventative and early intervention strategies (Chanen et al., 2007; Kaess et al., 2014).

Adolescence is a unique period of enhanced plasticity, marked by heightened risk-taking behaviors and impulsive decisions that can be adaptive (Spear, 2000), but can undermine survival and have adverse long-term consequences (e.g., risky sexual behavior, substance use, see Shulman et al., 2016 for review). Contemporary models (Luna & Wright, 2016; Steinberg, 2010) conceptualize adolescent impulsive behavior as a normative peak in reward-driven and affective behaviors, which are adaptive for specializing the neurobiological pathways required for adult-like levels of cognitive and affective functioning (Larsen & Luna, 2018a; Luciana et al., 2012; Luna et al., 2015; Spear, 2000). Specifically these models propose a relative predominance of affective/reward systems over cognitive control systems (Shulman et al., 2016), biasing adolescent decision-making toward rewarding stimuli. Thus adolescence represents a period in which normative peaks in impulsivity and affective processing *already* undermine decision-making (Larsen & Luna, 2018), and aberrant developmental trajectories may contribute to the emergence of major psychopathologies during this period (Larsen & Luna, 2018; Paus et al., 2008).

To date, the majority of experimental studies concerning decision-making in BPD have been conducted in adults (a recent meta-analysis reported mean age of 27 - 30 years (Jeung et al., 2016; but see also Tay et al., 2017), and have pointed to altered valuation of expected outcomes (Paret et al., 2017) predominantly assessed using delay discounting (Barker et al., 2015; Berenson et al., 2016; Coffey et al., 2011; Ka et al., 2008a; Krause-Utz et al., 2016a; Lawrence et al., 2010; Lempert et al., 2019; Maraz et al., 2016), reversal learning (Berlin et al., 2005; Paret et al., 2016), or the Iowa Gambling Task (IGT; Black et al., 2009; Cackowski et al., 2014; LeGris et al., 2012; Minzenberg et al., 2008; Schuermann et al., 2011; Haaland et al., 2007), although findings on the latter are mixed (Gorlyn et al., 2013; McCloskey et al., 2009; Paret et al., 2017). Increased trait impulsivity (Barker et al., 2015; Bornovalova et al., 2005; Lawrence et al., 2010; Skodol et al., 2002) has been shown to contribute to increased discounting rates in BPD (Coffey et al., 2011; Voelker et al., 2008; Krause-Utz et al., 2016; Lawrence et al., 2010), and variability in decision-making in healthy individuals (Franken et al., 2008; Penolazzi et al., 2012; Raio et al., 2020), including risk-taking behavior in adolescents (Romer et al., 2009). Increased levels of emotional dysregulation in BPD (e.g., difficulties in emotion regulation DERS; Gratz & Roemer, 2004), particularly with regard to lacking access to emotion regulation strategies and impulse control difficulties (Ibraheim et al., 2017), may lead to exacerbated impulsivity, particularly in the context of negative affect (Chapman et al., 2008; Hallquist et al., 2018; Koenigsberg et al., 2001; Krause-Utz et al., 2016a; J. R. Peters et al., 2013; Scott et al., 2014; Sebastian et al., 2013; Turner et al., 2017). Although developmentally appropriate assessment tools (including self-report, parent report, interview, and clinician report) have recently been validated to assess dysfunction in childhood and adolescence (Fonagy et al., 2015; Carla Sharp, Ha, et al., 2012; Carla Sharp & Fonagy, 2015), few experimental tasks that are sensitive to the pathological processes of BPD have been optimized for developmental populations. Here, we characterize how individual differences across two core features of personality psychopathology, impulsivity and emotional dysregulation, affect choice behavior during an interpersonal, competitive, decision-making task in adolescents with BPD.

Neuroeconomic games that probe decision-making in ecologically valid (often interpersonal) contexts (Hasler, 2012; Jeung et al., 2016; King-Casas & Chiu, 2012; Kishida et al., 2010; Montague, 2007; Robson et al., 2020; Sharp, 2012; Sharp et al., 2012) have revealed deficits in cooperative and trust behaviors in BPD, specifically related to perception of social norms and risk as compared to non-social risk (e.g., decisions involving fixed gambles; Franzen et al., 2011; Henco et al., 2020; King-casas et al., 2008; Preuss et al., 2016; Seres et al., 2009; Sharp, 2012; Unoka et al., 2009). Similarly, tasks assessing so-called ‘theory of mind’ processes (also known as ‘mentalizing’) have shown that adolescents with BPD show a tendency to ‘hypermentalize’ when inferring the intentions or mental states of others (Arntz et al., 2009; Henco et al., 2020; Sharp et al., 2011; Vaskinn et al., 2015; Zabihzadeh et al., 2017). Although these studies suggest changes in mentalization and cooperative decision-making processes in BPD, behavior during competitive interactions has not been investigated. Here, we explore how decisions made in a dynamic strategic context, which demands ongoing predictions of opponent’s choice behavior, differ in adolescents with BPD and as a function of impulsivity and emotional dysregulation.

During a game of rock-paper-scissors, each player’s actions and their associated outcomes change dynamically based on their opponent’s actions (Parr et al., 2019; Seo & Lee, 2008; von Neumann & Morgenstern, 1944; Vickery et al., 2011). On the one hand, it may be advantageous to adopt a mixed-strategy by choosing each of the three actions with equal frequency, but unpredictably from trial-to-trial. If both players do so, they approach the Nash equilibrium, and there is no incentive to deviate from this strategy unilaterally as departures could be exploited by their opponent (Fudenberg & Tirole, 1991; Nash, 1950). On the other hand, if one’s opponent deviates from the Nash equilibrium (e.g., by displaying preferences for one action or the other and/or serial dependence in choice patterns), one should be prepared to exploit these predictabilities, engaging mentalization processes involved in inferring the actions and mental states of others (Hampton et al., 2008; Vickery et al., 2015). Generally, individuals approach the Nash equilibrium, however, systematic deviations consistently emerge in normative studies of strategic decision-making in humans (Mikulić & Dorris, 2008; Parr et al., 2019; Vickery et al., 2011) and non-human primates (Barraclough et al., 2004; Lee et al., 2004; Thevarajah et al., 2009). Specifically, the win-stay/lose-shift (WSLS) bias emerges, consistent with the use of reinforcement learning (RL) processes (Cohen & Ranganath, 2007; Hampton et al., 2008), in which individuals are more likely to repeat previously successful actions (rewarded) and switch away from previously unsuccessful (unrewarded or punished) actions (Barraclough et al., 2004; Thevarajah et al., 2009; Vickery et al., 2011). Importantly, this paradigm is sensitive to several domains of dysfunction in BPD; 1. Impulsivity, given the inherently rapid and dynamic nature of the task that involves learning and updating the value of available actions (action-outcome contingencies) based on reinforcement history (Lee & Seo, 2007; Vickery et al., 2011) and translating this information into goal-directed actions; 2. Emotional dysregulation, given that affective sources of information are used to update value representations (Neville et al., 2021; Paulus et al., 2005), and 3. Interpersonal dysfunction, given that our task may recruit mentalization processes required to predict and draw inferences about the behavior of the opponent.

Based on the core clinical features, we hypothesized that individuals with BPD would show changes in mixed-strategy performance compared to age-matched controls. Pathological processes could affect strategic choice behavior in several ways. Impaired affective and RL processes could contribute to strategic choice behavior through outcome evaluation processes in the following ways; 1. Blunted reward prediction error (RPE) and/or affective signaling (Hüpen et al., 2020; Neville et al., 2021) that could lead to insensitivity to changing reward contingencies and/or a deficit updating the value of actions on a trial-by-trial basis, which in this context, may actually result in *fewer* choice biases (and therefore, potentially enhanced performance as one may better evade exploitation by the opponent); and/or 2. Exacerbated RPE and/or affective signals could lead to *amplified* biases (and therefore, potentially diminished performance as the opponent would exploit these biases as they emerge), either of which may be more prominent for either positive (WS) and/or negative (LS) outcomes (Neville et al., 2021; Paret et al., 2016; Schuermann et al., 2011). Relating to the mentalization findings in BPD, hypermentalization could contribute to strategic choice behavior through; 1. Exaggerated vigilance to social information that may result in enhanced prediction of the opponent’s behavior (and therefore improved task performance) and/or 2. Misattribution of opponent’s intentions/actions and/or overinterpretation of one’s own emotional responses (Henco et al., 2020; Sharp et al., 2011), resulting in diminished prediction of the opponent’s behavior (and therefore worse task performance). Given that this is a first-of-its kind study, we tested the possibility that BPD pathology could lead to either a detriment or a benefit to task performance. We further hypothesized that group differences would be more pronounced in individuals with higher levels of impulsivity in either group, measured using the BIS (Patton et al., 1995), and emotion dysregulation in the BPD group, measured using the DERS (Gratz & Roemer, 2004). Given evidence suggesting that emotion dysregulation and negative affect may partially exacerbate impulsivity in BPD (Neville et al., 2021; Sharp et al., 2011), we tested whether DERS mediated the relationship between impulsivity and choice behavior in BPD. Last, we conducted exploratory analyses to assess whether individual differences in self-report strategies contribute to the observed relationships between impulsivity (both groups) and difficulties in emotional regulation (BPD group) and choice behavior.

## Materials and Methods

### Participants

Female adolescents with BPD and age- and sex- matched control adolescents participated in an experiment that examined choice patterns during the mixed-strategy game, Matching Pennies (a two-choice variant of Rock-Paper-Scissors). Twenty-eight female adolescents (mean age: 16.3 years, ± 1.3, range: 14-18) who met criteria for BPD (assessed by the Structured Clinical Interview for DSM-5 Personality Disorders (SCID-PD; Pfohl et al., 1997) were recruited from the Dialectical Behavioral Therapy (DBT) group at Kingston Health Sciences Center outpatient mood and anxiety clinic at Hotel Dieu Hospital by co-author SKK. One BPD participant was excluded due to poor performance that indicated a lack of understanding of the task procedures (various performance measures including overall reward rate falling ≥ 3 SD from the group mean). Final analyses included twenty-seven female participants with BPD (mean age: 16.4 years, ± 1.3, range: 14-18) and twenty-seven female control participants (mean age: 15.8 years, ± 1.6, range: 14-18) recruited from the community in Kingston, Ontario. This study was approved by the Queen’s University Human Research Ethics Board and was in accordance with the *Canadian Tri-Council Policy Statement on Ethical Conduct for Research Involving Humans* and the principles of the *Declaration of Helsinki*. All participants gave informed consent and were compensated for their time. Both BPD and control participants completed an evaluation of impulsivity (Barratt Impulsiveness Scale (BIS-11; Patton et al., 1995)) as well as a post-game questionnaire (developed for the current study, see Methods section below) assessing how participants approached the strategic game (referred to as “Strategic Assessment”). Mean scores are shown in Table 1. BPD patients underwent an evaluation of borderline-typical symptomology (Borderline Symptom List (BSL-23); Bohus et al., 2007) and emotional dysregulation (Difficulties in Emotional Regulation Scale (DERS); Gratz & Roemer, 2004). Mean clinical scores for the BPD group are shown in Table 2. All BPD participants remained on their regular medication regimes throughout the duration of the study (medication information is shown in Table 2). Groups did not differ in terms of age (*BPD Mean* = 16.4 years, *SD* = 1.28, *CTRL Mean* = 15.8 years, *SD* = 1.55, *t* (52) = 1.44, *Cohen’s d* = .39, 95% CI [−.21, 1.32], *p* = .15), or handedness (*BPD*: 24 righthanded; *CTRL*: 26 righthanded; *t* (52) = .85, *Cohen’s d* = .23, 95% CI [−.15, .37], *p* = .39).

**Table 1.**
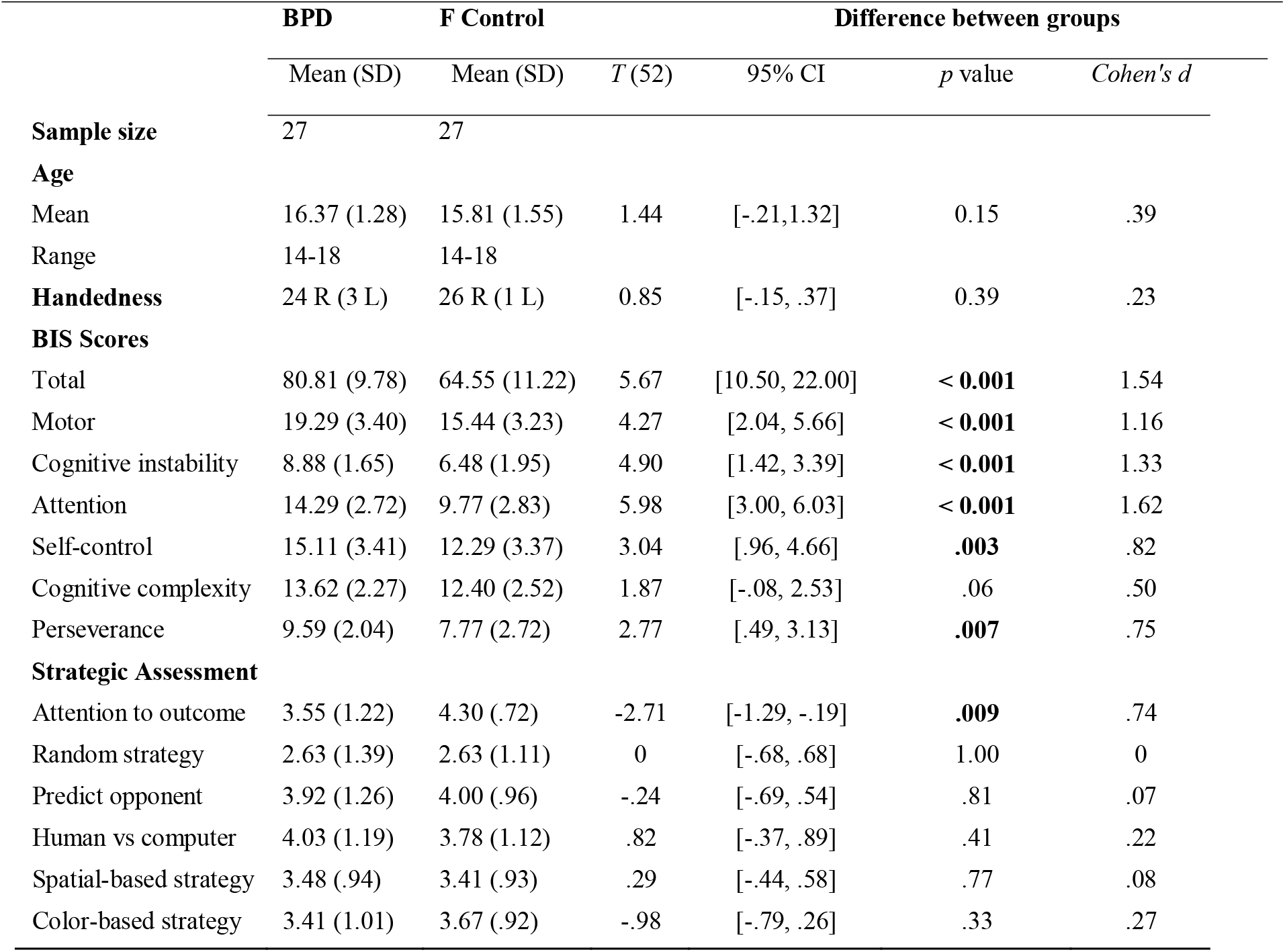
Descriptive Statistics of Participants. Data are reported as means (SD) unless otherwise indicated. BPD, borderline personality disorder. F, female. BIS, Barratt Impulsiveness Scale (BIS-11). P values are Bonferroni adjusted. Independent samples t-test.

**Table 2.** Clinical Scores and Medication Information for BPD Participants. Questionnaire data are reported as means (SD), while ADHD medication data are reported as number (percentage) of participants on that class of medication. BSL, Borderline Symptom List. DERS, Difficulties in Emotional Regulation Scale. Stimulant refers to Concerta, Methlyphenidate, Ritalin or Biphentin. Non-Stimulant refers to Intuniv or Straterra. Note that percentage of participants is based on the number of participants we have medication data for (N=25).

|  | **BPD** |
| --- | --- |
|  | Mean (SD) |
| **BSL<sup>a</sup>** | 53.80 (23.75) |
| **DERS<sup>b</sup>** |  |
| Total | 127.73 (23.58) |
| Non-acceptance | 21 (6.14) |
| Goals | 20.23 (4.16) |
| Impulse control | 21.58 (6.27) |
| Awareness | 20.19 (5.00) |
| Strategies | 28.54 (6.40) |
| Clarity | 16.19 (4.46) |
| **Medication (Class)<sup>c</sup>** | N (% of participants) |
| Stimulant | 6 (24) |
| Non-Stimulant | 3 (12) |

<sup>a</sup> Data not available for two patients (N=25)
<sup>b</sup> Data not available for one patient (N=26)
<sup>c</sup> Data not available for two patients (N=25)

### Task Procedures

#### Strategic Decision-Making Task

Participants competed in a color-based version of Matching Pennies against a dynamic computer opponent that exploited predictabilities in player choice patterns (Fig. 1A). Participants played the role of the *matcher*, while the computer opponent played the role of the *non-matcher* - if both players chose the same colored target, the participant won $0.10; otherwise, the participant lost $0.10 (Fig. 1B; players were endowed with 30 credits at the beginning of the session). Briefly, the competitive algorithm employed by our computer opponent was based on algorithm 2 in Barraclough et al., 2004, and performed a statistical analysis of participants’ historical sequence of choices (including both leftward/rightward target and red/green target) and associated payoffs (rewarded or unrewarded) to uncover systematic biases (Barraclough et al., 2004 and Parr et al., 2019). Participants were informed of the rules of the game, were aware that they were playing a strategic game against a dynamic, competitive, computer opponent, and were instructed to win as much money as possible. If participants approached the Nash equilibrium (e.g., by successfully evading exploitation by the opponent and/or randomizing over choice patterns), they would win approximately 50% of trials.

**Figure 1.**
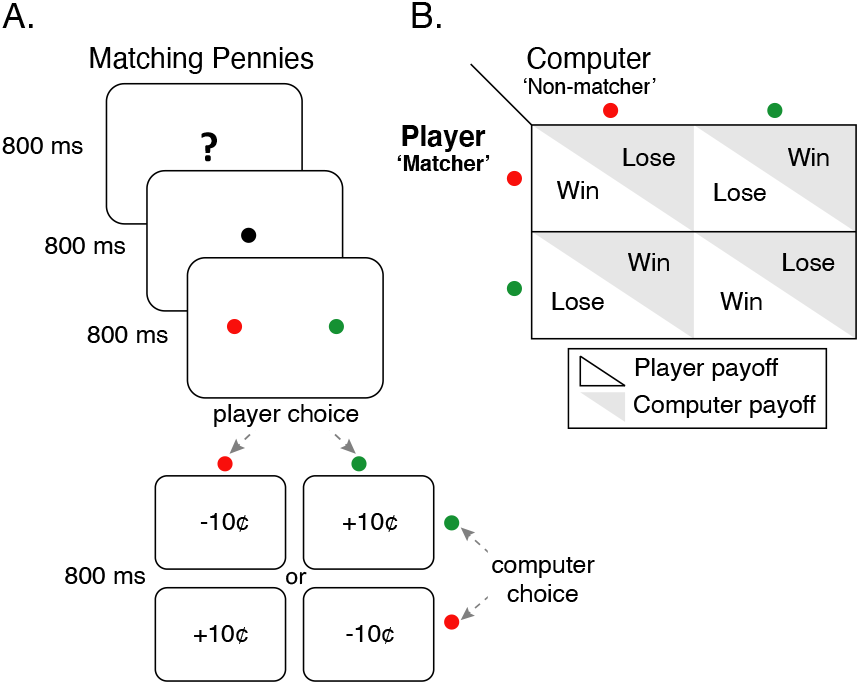
Matching Pennies. (A) Participants were informed that they were competing in Matching Pennies against a dynamic computer opponent that analyzed their behavior and exploited predictabilities in their response patterns (see algorithm 2 in Lee et al., 2004 and Parr et al., 2019). Participants played the role of the ‘matcher’ while the computer opponent played the ‘non-matcher’; if both players chose the same colored target, participants received a ‘10¢’ reward for that trial. Otherwise, participants lost ‘10¢’ for that trial. They were instructed to win as much money as possible. (B) Payoff matrix for each player.

Participants were informed that monetary compensation for the current study was dependent on task performance (cumulative amount of reward earned) during Matching Pennies. However, independent of task performance, each participant was compensated with a $30 gift card.

#### Experimental Design and Timing

Participants completed four runs (each consisting of 150 trials for a total of 600 trials) of Matching Pennies (Fig. 1A). Each trial was 3200 ms in duration and started with 800 ms of a task identification period followed by 800 ms of a fixation period. Next, two visual targets, one green and one red, appeared for 800 ms at an eccentricity of 6.5° to the left and the right of the fixation point, during which participants indicated their choice of target with a saccadic eye movement. Finally, the outcome of each trial (monetary reward) was revealed during an 800 ms period. Participants were presented with a 20 second long fixation period at the beginning and end of each run. Each run was 8 minutes and 40 seconds in duration, and the total session time was approximately 60 minutes (including breaks between runs and approximately 15 minutes for questionnaires). Each participant completed 20 practice trials at the beginning of the session.

Importantly, the location of the colored targets was pseudo-randomized, appearing on each side of the screen 50% of the time. Participants were instructed to maintain fixation prior to the appearance of the targets. Saccades made prior to the appearance of the targets were considered anticipatory trials and those made >800ms following the appearance of the targets were considered non-response trials.

#### Recording and Apparatus

Monocular eye position data was recorded at 500 Hz using the EyeLink 1000 Version 5.1 table mounted eye-tracking device (SR-Research Ltd., Mississauga, Ontario, Canada). The monitor, infrared illuminator, and camera were positioned 60 cm away from central gaze, and the right eye was recorded. Participants were situated in a mounted chin rest, stabilizing the head and limiting motion during each trial. All visual stimuli were presented and behavioral responses acquired using custom MATLAB v7.9 programs (The Mathworks Inc., Natick, MA) and Psychophysics Toolbox v3 (Brainard, 1996; Pelli, 1996) running on a PC. Visual stimuli were presented on an adjustable 17-inch LCD monitor at a screen resolution of 1280×1024 pixels that had a refresh rate of 60 Hz. At the beginning of each run, eye position was calibrated using a five-point calibration routine set to within 1° of the visual target. Participants indicated their choices by making a saccadic eye movement that corresponded to the location of the desired visual target. Saccades were recorded if amplitudes were > 7.5°.

#### Performance Variables

The following behavioral variables were examined: (1) the probability of reward (p(rew)), which served as a proxy for overall task performance. We also measured the extent to which participants’ choices depended on a history of previous choices and outcomes by calculating (2) the probability of using the win-stay-lose-shift strategy (p(wsls)), choosing the same target after a reward in the previous trial (win-stay) and/or switching to the opposite target following a loss in the previous trial (lose-shift).

To quantify the degree of randomness in participants’ choice patterns (e.g., the degree of randomness in participants choice patterns), we calculated (3) entropy, termed choice entropy (Cover & Thomas, 1991). We also calculated (4) entropy using the choice sequence of the two players (which is equivalent to using outcome), termed choice-outcome entropy (see Lee et al., 2004 for details).

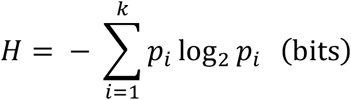

Calculating the entropy based on the participant’s choice (choice entropy) in three successive trials, there are a total number of eight possible outcomes (*k* = 2^3^ = 8), and the maximum entropy is 3 bits. When entropy is calculated based on the participant’s choice sequence in three successive trials, as well as the computer opponent’s choice sequence in two prior trials (choice-outcome entropy), there are a total number of 32 possible outcomes (k = 2^5^ = 32), and the maximum entropy is 5 bits. Entropy was log transformed for all analyses.

Both p(wsls) and entropy variables were computed separately for both color and spatial domains (i.e., if participants chose the righthand target following a rewarded outcome on the right, this would be considered spatial wsls bias).

#### Response Time Variables

We additionally recorded saccadic reaction times (SRT), and calculated the coefficient of variation in SRT (CV; *CV* = SD / Mean*100), the percentage of anticipatory trials, and the percentage of non-response trials.

### Assessments

#### Barratt Impulsiveness Scale (BIS-11)

In both groups, we investigated whether individual variation in impulsivity corresponded to strategic choice behaviors. The Barratt Impulsiveness Scale (BIS-11; Mean scores for each subscale are shown in Table 1) is a well validated tool (Barratt 1985; Patton et al., 1995; Stanford et al., 2009), and six first-order factors (sub-traits) have been identified (Patton et al., 1995) that reflect impulsivity across the following domains: 1. Attention (e.g., “I concentrate easy”, “I am a steady thinker”, “I am restless in lectures”); 2. Motor (e.g., “I do things without thinking”, “I act on impulse/on the spur of the moment”, “I buy things on impulse”); 3. Self-control (e.g., “I plan tasks carefully”, “I am self-controlled”, “I say things without thinking”); 4. Cognitive complexity (e.g., “I like to think about complex problems”, “I am more interested in the present than the future”, “I like puzzles”); 5. Perseverance (e.g., “I can only think about one thing at a time”, “I am future oriented”, “I change jobs”); and 6. Cognitive instability (e.g., “I have racing thoughts”, “I change hobbies”, “I often have extraneous thoughts when thinking”; Patton et al., 1995; Stanford et al., 2009). Higher scores on each scale indicate greater impulsivity (mean scores for each subscale are shown in Table 1).

#### Difficulties in Emotional Regulation (DERS)

In the BPD group, we investigated whether choice patterns changed as a function of emotional dysregulation. The Difficulties in Emotional Regulation Scale (DERS; Gratz & Roemer, 2004; mean scores for each subscale are shown in Table 2) measures functioning across several domains: 1. Nonacceptance of emotional responses (non-acceptance; e.g., “When I’m upset, I become irritated with myself for feeling that way”); 2. Difficulties engaging in goal-directed behavior (goals; e.g., “When I’m upset, I have difficulty thinking about anything else”); 3. Impulse control difficulties (impulse control; e.g., “When I’m upset, I have difficulty controlling my behaviors”); 4. Lack of emotional awareness (awareness; e.g., “When I’m upset, I (do not) acknowledge my emotions”); 5. Limited access to emotion regulation strategies (strategies; e.g., “When I’m upset, I start to feel very bad about myself”); and 6. Lack of emotional clarity (clarity; e.g., “I have difficulty making sense out of my feelings”; Gratz & Roemer, 2004). Higher scores indicate greater emotional dysregulation (mean scores for each subscale are shown in Table 2). Importantly, DERS data were not collected in control participants, thus we do not present group comparisons for this measure.

Impulsivity (Stanford et al., 2009) and emotional dysregulation (Gratz & Roemer, 2004) are multifaceted constructs, and differential patterns of associations among impulsivity and DERS subscales have been associated with distinct clinically relevant outcomes (Gratz & Roemer, 2004; Krause-Utz et al., 2016; Paret et al., 2016), including externalizing and internalizing characteristics in adolescents (Neumann et al., 2010). Because our study was novel and exploratory in nature, we report means across each subscale and examine the contribution of each subscale to strategic choice behavior.

#### Strategic Assessment

In both groups, we administered a post-game questionnaire that was developed for the current study to assess whether the way in which participants approached the mixed-strategy game affected strategic choice patterns. The questionnaire was comprised of the following items: 1. Attention to outcome: “I paid attention to whether I was rewarded or not, and changed by strategy when I wasn’t being rewarded (or when I was)”; 2. Spatial strategy: “I decided which target to choose before the targets appeared on the screen (I chose based on left or right side of the screen)”; 3. Color strategy: “I decided which target to choose based on the color of the targets”; 4. Random strategy: “I found that it was easy to choose randomly”; 5. Exploit opponent: “I tried to predict what my opponent would do”; and 6. Human versus computer: “I think I would play differently if I was playing against a human opponent as opposed to a computer”. Each item was rated on a 5-point scale with 1 representing “Strongly Disagree” and 5 representing “Strongly Agree”, and we assessed the relationship between each item and choice behavior (mean scores for each item are shown in Table 1).

### Data Analysis

Statistical analyses were carried out with custom MatLab programs version 9.3 (The MathWorks Inc., MA) and R version 3.5.2 via RStudio version 1.2.1 (R Core Team, 2017; RStudio Team, 2020). All continuous variables were z-scored prior to statistical analyses and z-scores are shown in scatterplots to allow for visualization across different assessments.

#### Group Differences in Task Performance

We first evaluated whether there were significant differences between groups on the dependent variables affecting Matching Pennies performance, including 1) reward rate; 2) win-stay, lose-shift biases; 3) choice entropy; and 4) choice-outcome entropy. The latter three measures were calculated for both the spatial and color domains. Performance data were considered significant at *p* < .01 to account for multiple comparisons (Bonferroni correction: 0.05/4 performance variables, corrected alpha = .01).

Next, we evaluated differences between groups on the dependent variables affecting response times, including 1) median SRT; 2) CV; 3) % of anticipatory trials; and 4) % of non-response trials. Data were considered significant at *p* < .01 to account for multiple comparisons (Bonferroni correction: 0.05/4 behavioral variables, corrected alpha = .01).

To explore whether any group differences were being driven by BPD participants who were taking stimulant medication (to treat symptoms of attention deficit hyperactivity disorder (ADHD)), we further tested for differences between BPD participants prescribed stimulants for comorbid ADHD (n=6), and those not prescribed stimulants (n=19; see Table 2).

In all cases, two samples t-tests were conducted to investigate between-group differences on individual dependent variables where there was homogeneity of variances between groups (using Shapiro-Wilks tests), otherwise, Mann-Whitney U tests were conducted. Effect sizes are reported for all comparisons.

#### Associations with BIS and DERS scores

We next conducted linear regression models to examine the relationship between impulsivity (each subscale of the BIS) and choice behaviors, and whether this differed as a function of group (BPD versus control). We first tested for assessment by group interactions, and interaction terms were removed from the final models when there were no significant interaction terms. In the BPD group, we examined the relationship between emotional dysregulation (DERS) and choice behaviors.

#### Associations with Strategic Assessment (Exploratory Analyses)

We conducted linear regression models to explore the relationship between self-reported strategies (Strategic Assessment, see methods) and choice behaviors, and whether this differed as a function of group (BPD versus control). As above, we first tested for assessment by group interactions, and interaction terms were removed from the final models when there were no significant interaction terms.

For all above tests, data were considered significant at *p* < .01. Correlation matrices (ggcorrplot, R) were computed for visualization of correlations among variables in each group.

To explore whether any associations were being driven by BPD participants who were taking stimulant medication (n=6) to treat symptoms of attention deficit hyperactivity disorder (ADHD, n=6), we repeated these models testing for assessment (BIS, Strategic Assessment & DERS) by stimulant interactions in the BPD group, and interaction terms were removed from the final models when there were no significant interaction terms.

#### Mediation Analyses

We performed mediation analyses to test the hypothesis that DERS mediated the effects of impulsivity on choice behavior in BPD. The linear regression models employed in the mediation analyses were implemented as in the previous analyses with the same covariance structure. Significance values for indirect effects were obtained using 10000 draws in a bootstrap procedure (mediation package in R; Tingley et al., 2014).

Finally, we conducted exploratory mediation analyses to explore whether DERS may partially account for the relationship between self-report strategies and choice behavior in BPD.

## Results

### Group Differences in Task Performance

#### Matching Pennies Performance

BPD and control participants were comparable on the majority of performance variables (Table 3), with the exception of the probability of win-stay, lose-shift in the color domain, which was lower in the BPD group (Fig. 2 & Table 3; *M* = .51, *SD* = .02) as compared to the control group (*M* = .54, *SD* = .03; *t (52)* = −2.14, *Cohen’s d* = .58, *p* = .03, *p_Bonferroni_* = .12), although this failed to survive multiple comparison corrections despite the moderate effect size. Overall reward rate did not differ among groups (Fig. 3A, *p* > .05), and performance variables did not differ among BPD participants as a function of stimulant medication (all *p* > .05).

**Table 3.** Performance on Matching Pennies Task. Data are reported as means (SD) unless otherwise indicated. BPD, borderline personality disorder. F, female. P (win), probability of winning on average. P(WSLS), probability of win-stay, lose-shift strategy. Choice entropy, entropy in choice sequence in three successive trials and the choice in the next trial. Choice-outcome entropy, entropy in choice sequence of two players in two successive trials and the subject's choice in the next trial. RT, reaction time. ms, milliseconds. denotes Mann-Whitney U Test, in which case the test statistic is rank, effect size is ‘r’, and medians are reported. All other results were based on T-Tests, in which case the test statistic is ‘T’, effect size is Cohen’s D, and means are reported.

| Measure | BPD | F control | Test Statistic | p value | $p_{Bonferroni}$ | 95% CI | Effect Size |
| --- | --- | --- | --- | --- | --- | --- | --- |
|  | Mean (SD) | Mean (SD) |  |  |  |  |  |
| p (rew) | .47 (.03) | .47 (.02) | -.08 | .93 | 1.00 | -1.75, 1.61 | .02 |
| **Spatial Biases** |  |  |  |  |  |  |  |
| p (wsls) | .55 (.06) | .56 (.05) | -.40 | .68 | 1.00 | -.03, .02 | .11 |
| choice entropy | 2.96 (.07) | 2.94 (.06) | 444 $\psi$ | .17 | 1.00 | -.007, .03 | .12 |
| choice-outcome entropy | 4.81 (.13) | 4.81 (.09) | 335 $\psi$ | .61 | 1.00 | -.05, .03 | .04 |
| **Color Biases** |  |  |  |  |  |  |  |
| p (wsls) | .51 (.02) | .54 (.03) | -2.14 | **.03** | .21 | -.03, -.001 | **.58** |
| choice entropy | 2.99 (.01) | 2.99 (.02) | 438 $\psi$ | .20 | 1.00 | -.001, .009 | .11 |
| choice-outcome entropy | 4.95 (.03) | 4.93 (.03) | 458 $\psi$ | .10 | .70 | -.003, .03 | .16 |
| **Response variables** |  |  |  |  |  |  |  |
| Median RT (ms) | 258 (5.53) | 266 (6.46) | 333.5 $\psi$ | .59 | 1.00 | -22.00, 11.99 | .03 |
| CV (ms) | 270.25<br>(207.37) | 135.56<br>(141.83) | 2.27 | **.02** | .08 | 1.46, 23.52 | **.61** |
| % anticipatory | 6.71 (3.63) | 3.32 (2.57) | 3.95 | **< .001** | **<.001** | 1.65, 5.15 | **1.07** |
| % non-response | 1.42 (1.91) | .46 (.56) | 2.49 | **.02** | **.07** | .18, .95 | **.68** |

**Figure 2.**
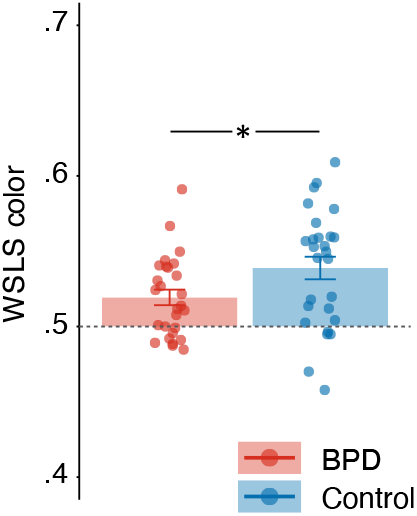
Group Differences in the Probability of Win-Stay, Lose-Shift (p(wsls)) in the color domain. Independent samples T-Test results are reported, and reported *p* value is uncorrected. * *p* < .05, ** *p* <.01, *** *p* < .001.

**Figure 3.**
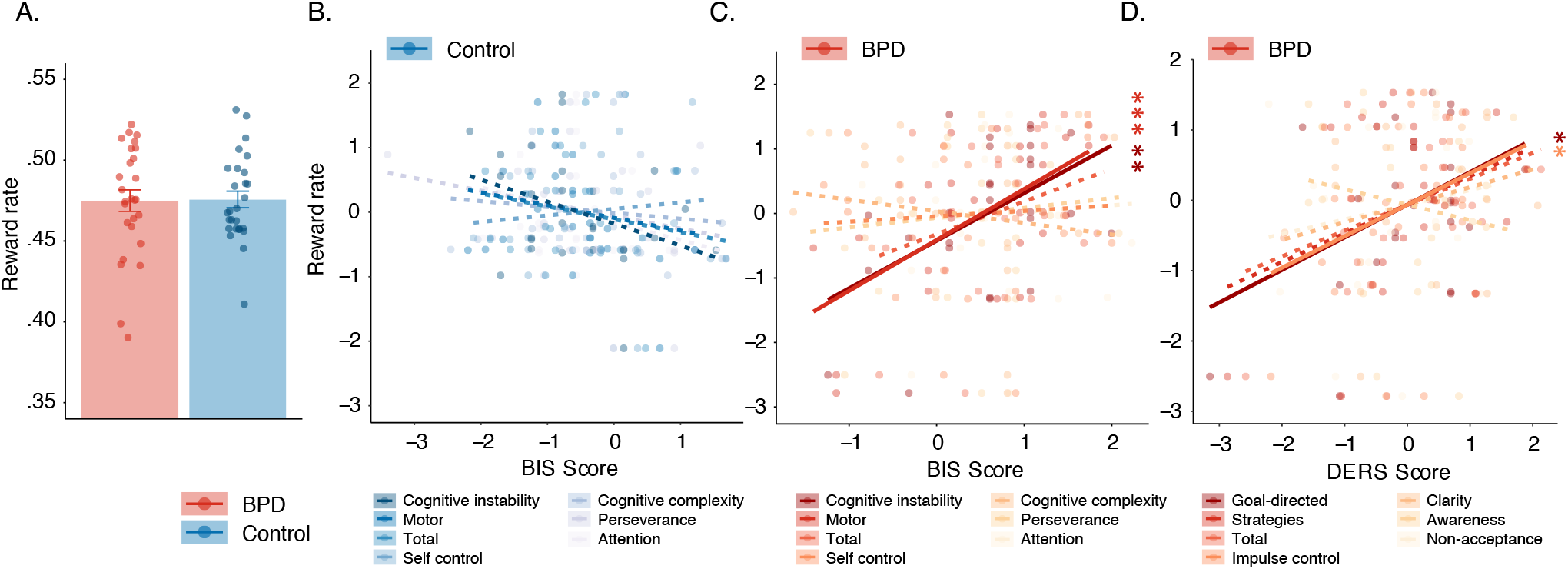
Reward Rate During Matching Pennies. (A) Group differences in reward rate. The relationship between reward rate and BIS impulsivity scores in (B) control and (C) BPD participants. (D) The relationship between reward rate and DERS emotion dysregulation scores in BPD participants. In A, independent samples T-Test results are reported, in B-D, z-scores are shown to allow for visualization across different subscales, linear effects results are reported, and solid and dashed lines indicate significant and non-significant regressions, respectively, and reported *p* values in B & C are Bonferroni corrected, while D is uncorrected. * *p* < .05, ** *p* <.01, *** *p* < .001.

#### Response Time Variables

CV was significantly higher in the BPD group (Fig. 4A & Table 3; *M* = 270.25 ms, *SD* = 207.37 ms) as compared to the control group (Fig. 4A & Table 3; *M* = 135.56 ms, *SD* = 141.83 ms; *t (52)* = 2.78, *Cohen’s d* = .76, *p* = .007, *p_Bonferroni_* = .03). Additionally, the percentage of anticipatory trials was significantly higher in the BPD group (Fig. S1A & Table 3; *M* = 6.71%, *SD* = 3.63%) as compared to the control group (Fig. S1A & Table 3; *M* = 3.32%, *SD* =2.57%; *t (52)* = 3.95, *Cohen’s d* = 1.07, *p* < .001, *p_Bonferroni_* <.001). The percentage of non-response trials was higher in the BPD group (Table 3; *M* = 1.42%, *SD* = 1.91%) compared to the control group (Table 3; *M* = .46%, *SD* = .57%; *t (52)* = 2.49, *Cohen’s d* = .68, *p = .018, p_Bonferroni_* =.07), although this failed to reach criteria for multiple comparisons and non-response rates were low overall (< 2% in both groups, Table 3). Median SRTs did not differ among groups (*W* = 333.5, *r* = .03, *p_Bonferroni_* = 1.00), and response time variables did not differ among BPD participants as a function of stimulant medication (all *p* > .05). Because CV and the percentage of anticipatory decisions were so highly correlated (*Pearson’s R* = .92 across the whole sample), we show CV results in Fig. 4 and anticipatory decisions in Fig. S1.

**Figure 4.**
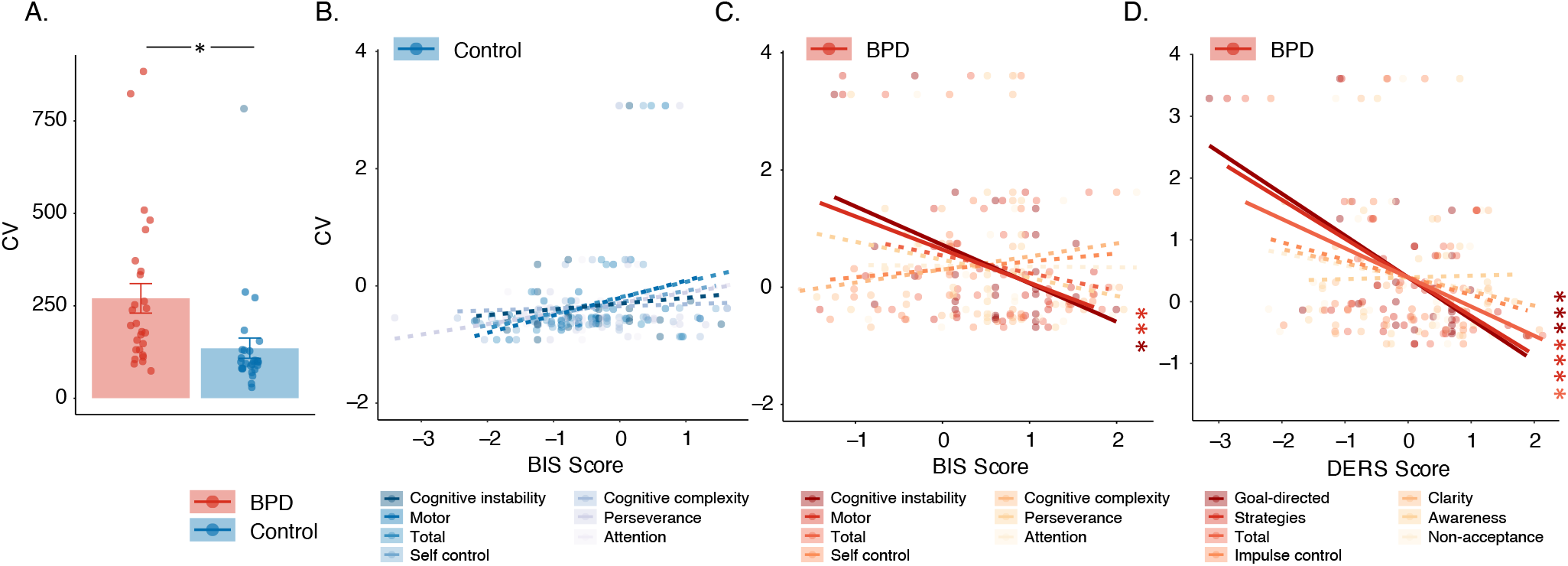
Coefficient of Variation in Saccadic Reaction Times (CV) During Matching Pennies. (A) Group differences in CV. The relationship between CV and BIS impulsivity scores in (B) control and (C) BPD participants. (D) The relationship between CV and DERS emotion dysregulation scores in BPD participants. In A, independent samples T-Test results are reported, in B-D, z-scores are shown to allow for visualization across different subscales, linear effects results are reported, and solid and dashed lines indicate significant and non-significant regressions, respectively, and reported *p* values are Bonferroni corrected. * *p* < .05, ** *p* <.01, *** *p* < .001.

### Relationship between Impulsivity and Matching Pennies Performance

Total BIS scores were higher in the BPD group (*M* = 80.81, *SD* = 9.78), compared to control participants (*M* = 64.55, *SD* = 11.22; *t* (52) = 5.67, *Cohen’s d* = 1.54, *p* < .001). Broken down into the individual subscales, individuals with BPD scored significantly higher on the cognitive instability, motor, attention, self-control, and perseverance subscales (all *p* < .05, Table 1), but did not differ from control participants in terms of the cognitive complexity subscale (Table 1).

#### BIS-Motor Subscale

We found a significant interaction between group and BIS-MO on overall reward rate (Fig. 3A; β = −.99, *t* = −3.47, *p* = .001*; p_Bonferroni_* = .004), and *post hoc* tests revealed a significant positive association between BIS-MO scores and reward rate in the BPD group (Fig. 3C; β = .79, *t* = 3.97, *p* < .001), but a non-significant negative association between BIS-MO scores and reward rate in the control group (Fig. 3B; β = −.20, *t* = −.97, *p* = .34). The main effect of BIS-MO on reward rate did not meet criteria for multiple comparisons (β = .32, *t* = 2.05, *p =* .05, *p_Bonferroni_* = .20).

We also found a significant interaction between group and BIS-MO on CV (β = .86, *t* = 3.10, *p* = .003, *p_Bonferroni_* = .01), and *post hoc* tests revealed a negative association between BIS-MO scores and CV in the BPD group (Fig. 4C; β = −.57, *t* = −2.58, *p* = .02), whereas a trend level positive association was observed in control participants (Fig. 4B; β =.30, *t* =1.77, *p* = .09). We did not observe a significant main effect of BIS-MO on CV (β = −.16, *t* =-1.04, *p* = .30, *p_Bonferroni_* = 1.00). Additionally, we found an interaction between group and BIS-MO on the percentage of anticipatory trials (β = .63, *t* = 2.30, *p* =.03, *p_Bonferroni_* = .12), though this failed to reach criteria for multiple comparisons. Nonetheless, exploratory *post hoc* tests revealed a trend level negative association between BIS-MO scores and anticipatory trials in the BPD group (Fig. S1C; β = −.42, *t* = −1.98, *p* = .06) and a non-significant positive association in the control participants (Fig. S1B; β = .21, *t* =1.24, *p* = .23). We did not observe a significant main effect of BIS-MO on anticipatory trials (β = −.12, *t* = −.87, *p* = .39, *p_Bonferroni_* = 1.00).

Finally, we observed a trend-level group by BIS-MO interaction on choice-outcome entropy (β = −.53, *t* = −2.38, *p* = .02, *p_Bonferroni_* = .08), and exploratory *post hoc* tests revealed a positive relationship between BIS-MO and choice-outcome entropy in the BPD group (β = .35, *t* = 2.20, *p* = .04) that was not evident in the control participants (β = −.18, *t* = −1.14, *p* = .26). We did not observe a significant main effect of BIS – MO on choice-outcome entropy (β = .10, *t* = .86, *p* = .39, *p_Bonferroni_* = 1.00).

#### BIS-Cognitive Instability Subscale

We found a significant interaction between group and BIS-CI on reward rate (β = −1.08, *t* = −3.48, *p* =.001, *p_Bonferroni_* = .004), and *post hoc* tests revealed a significant positive association between BIS-CI scores and reward rate in the BPD group (Fig. 3C; β = .74, *t* = 2.87, *p* = .01). On the other hand, control participants showed a trend toward a negative relationship between BIS-CI and reward rate (Fig. 3B; β = −.34, *t* = −1.88, *p* = .07). We did not observe a significant main effect of BIS-CI on reward rate (β = .11, *t* =.65, *p* = .52, *p_Bonferroni_* = 1.00).

We found an trend-level interaction between group and BIS-CI on CV (β = .75, *t* = 2.49, *p* = .02, *p_Bonferroni_* = .08), and exploratory *post hoc* tests revealed a negative association between BIS-CI scores and CV in the BPD group (Fig. 4C; β = −.66, *t* = −2.56, *p* = .02), but no significant association in control participants (Fig. 4B; β =.10, *t* =.59, *p* = .56). We did not observe a significant main effect of BIS-CI on CV (β = −.22, *t* =-1.40, *p* = .17, *p_Bonferroni_* = .68). Additionally, we observed a trend-level interaction between group and BIS-CI on the percentage of anticipatory trials (β = .63, *t* = 2.22, *p* =.03, *p_Bonferroni_* = .12), and *post hoc* tests revealed a significant negative association between BIS-CI scores and anticipatory trials in the BPD group (Fig. S1C; β = −.56, *t* = −2.28, *p* = .03) but no significant association in the control participants (Fig. S1B; β =.08, *t* =.49, *p* = .63). We did not observe a significant main effect of BIS-CI on anticipatory trials (β = −.19, *t* =-1.27, *p* = .21, *p_Bonferroni_* = .84).

Finally, we observed a trend-level group by BIS-CI interaction on choice-outcome entropy (β = −.57, *t* = −2.45, *p* = .02, *p_Bonferroni_* = .08), and exploratory *post hoc* tests revealed a significant negative relationship between BIS-CI and choice-outcome entropy in the control group (not shown; β = −.40, *t* = −2.93, *p* = .005), that was not evident in the BPD participants (not shown; β = .17, *t* = .83, *p* = .41). We did not observe a main effect of BIS – CI on choice-outcome entropy (β = −.17, *t* = −1.37, *p* = .19, *p_Bonferroni_* = .76).

We observed no significant associations between task performance variables and the BIS attention, self-control, cognitive complexity, and perseverance subscales (all *p* < .05, Fig. 7), and all relationships remained following the inclusion of age and stimulus medication (all *p* < .05).

**Figure 7.**
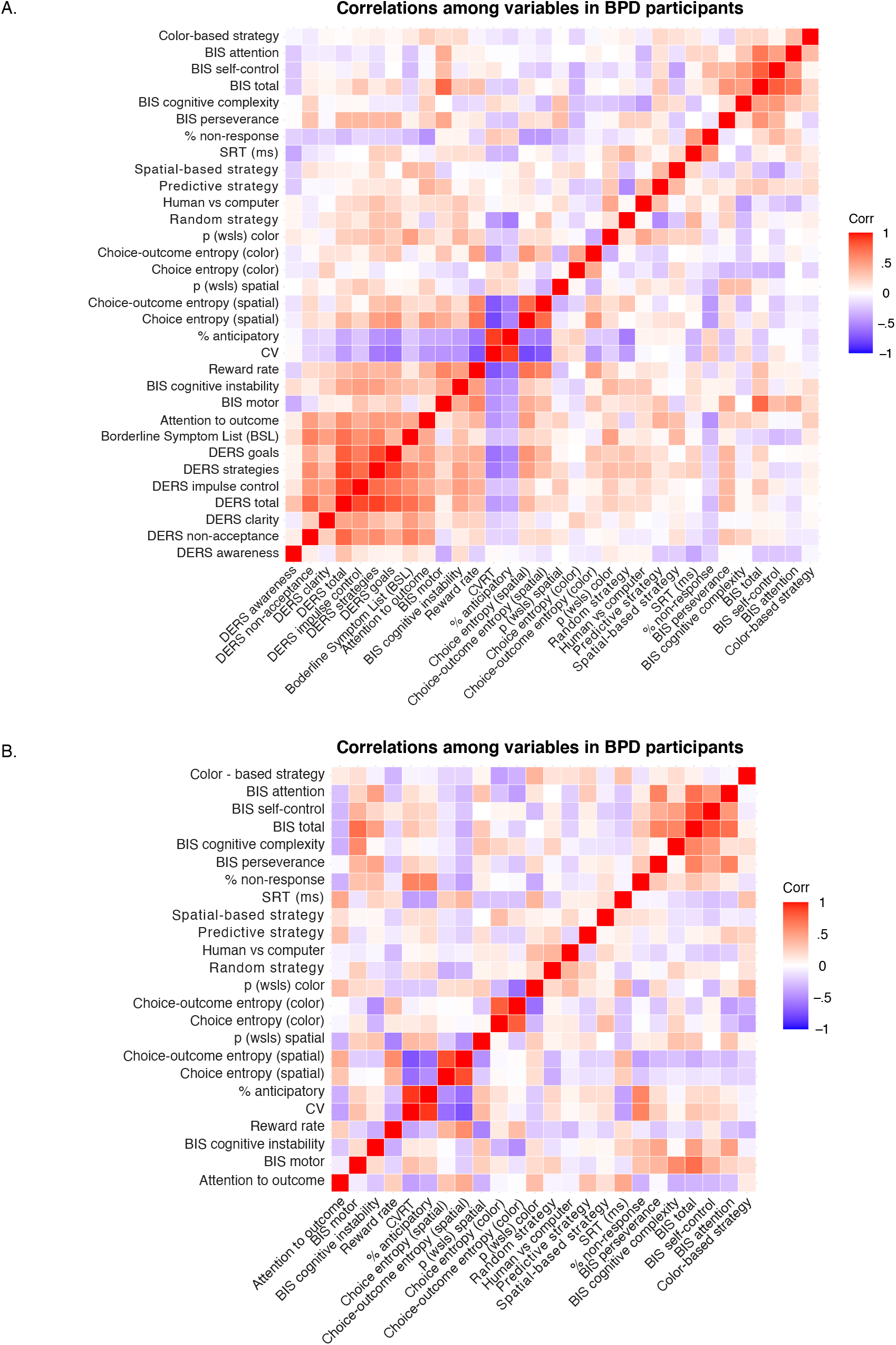
Correlation matrices showing relationships among behavioral variables and assessment data in (A) BPD and (B) control participants. ‘Hot’ and ‘cold’ colors reflect positive and negative relationships among variables, respectively.

### Relationship between DERS and Matching Pennies Choice Behavior in BPD

#### DERS Goals Subscale

In the BPD group, difficulties in engaging in goal directed behavior was significantly negatively associated with CV (Fig. 4D; β = −.68, *t* = −3.80, *p* < .001, *p_Bonferroni_* = .003) and the percentage of anticipatory trials (Fig. S1D; β = −.53, *t* = −2.98, *p* = .01, *p_Bonferroni_* = .04). Further, DERS goals was positively associated with reward rate (Fig. 3D; β = .46, *t* = 2.22, *p* =.04, *p_Bonferroni_* = .20), choice entropy (β = .30, *t* = 2.37, *p* = .02, *p_Bonferroni_* = .08) and choice-outcome entropy (not shown; β = .33, *t* = 2.27, *p* = .03, *p_Bonferroni_* = .12), although these relationships failed to meet criteria for multiple comparisons.

#### DERS Strategies Subscale

DERS strategies was significantly negatively associated with CV (Fig. 4D; β = −.63, *t* = −3.38, *p* = .002, *p_Bonferroni_* = .008), and we observed a trend level relationship with the percentage of anticipatory trials (Fig. S1D; β = −.47, *t* = −2.54, *p*=.02, *p_Bonferroni_* = .08).

#### DERS Impulse Control Subscale

Finally, we observed a trend-level positive relationship between DERS impulse and reward rate (Fig. 3D; β = .45, *t* = 2.13, *p* = .04, *p_Bonferroni_* = .16).

We observed no significant associations between task performance variables and the DERS clarity, non-acceptance, and awareness subscales (all *p* < .05, Fig. 7A), and all relationships remained following the inclusion of age and stimulus medication (all *p* < .05).

#### Correspondence between DERS & BIS

To understand the common mechanisms by which our assessment data affects choice patterns during matching pennies, we highlight critical associations here (but see Fig. 7 for correlation matrices of all variables). Importantly, BIS-CI was significantly associated with the majority of DERS subscales (Fig. 7A; with the exception of non-accept and awareness; all *p* <.05), and we found no other associations between DERS and BIS scores across any other subscale (Fig. 7A).

#### DERS Mediates the Effects of Impulsivity on Choice Behavior in BPD

We conducted mediation analyses to test whether difficulties in emotional regulation may partially account for the observed relationship between impulsivity and choice behavior in BPD. We tested for mediation where we saw significant associations between choice behavior and both BIS scores and DERS scores, and found that the relationship between BIS – CI and CV was significantly mediated by both DERS goals (Fig. 5; β = −.31, CI [−.73, .−02], *p* = .04) and DERS strategies (Fig. 5; β = −.30, CI [−.72, −.01], *p* = .04).

**Figure 5.**
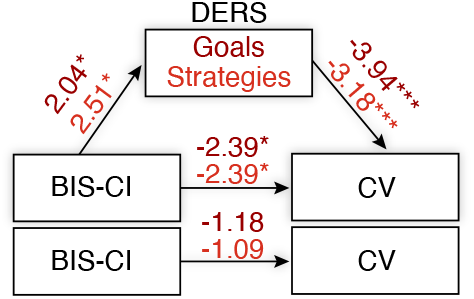
Difficulties in Emotional Regulation (DERS) Mediation Analyses in the BPD Group. DERS ‘Goals’ and ‘Strategies’ significantly mediated the relationship between impulsivity and coefficient of variation in saccadic reaction times (CV) in BPD. T-test results are reported, and significance values for indirect effects were obtained using 10000 draws in a bootstrap procedure (mediation package in R; Tingley et al., 2014).

### Exploratory Analyses: Relationship between Strategic Assessment and Matching Pennies

#### Attention to Outcome

As a group, individuals with BPD reported paying significantly less attention to outcome during the mixed-strategy game (Fig. 6A; *M* = 3.55, *SD* = 1.22) as compared to control participants (Fig 6A; *M* = 4.30, *SD* = .72; *t* (52) = −2.71, *Cohen’s d* = .74, *p* = .009). Otherwise, BPD and control participants did not differ in self-report strategies (Table 1).

**Figure 6.**
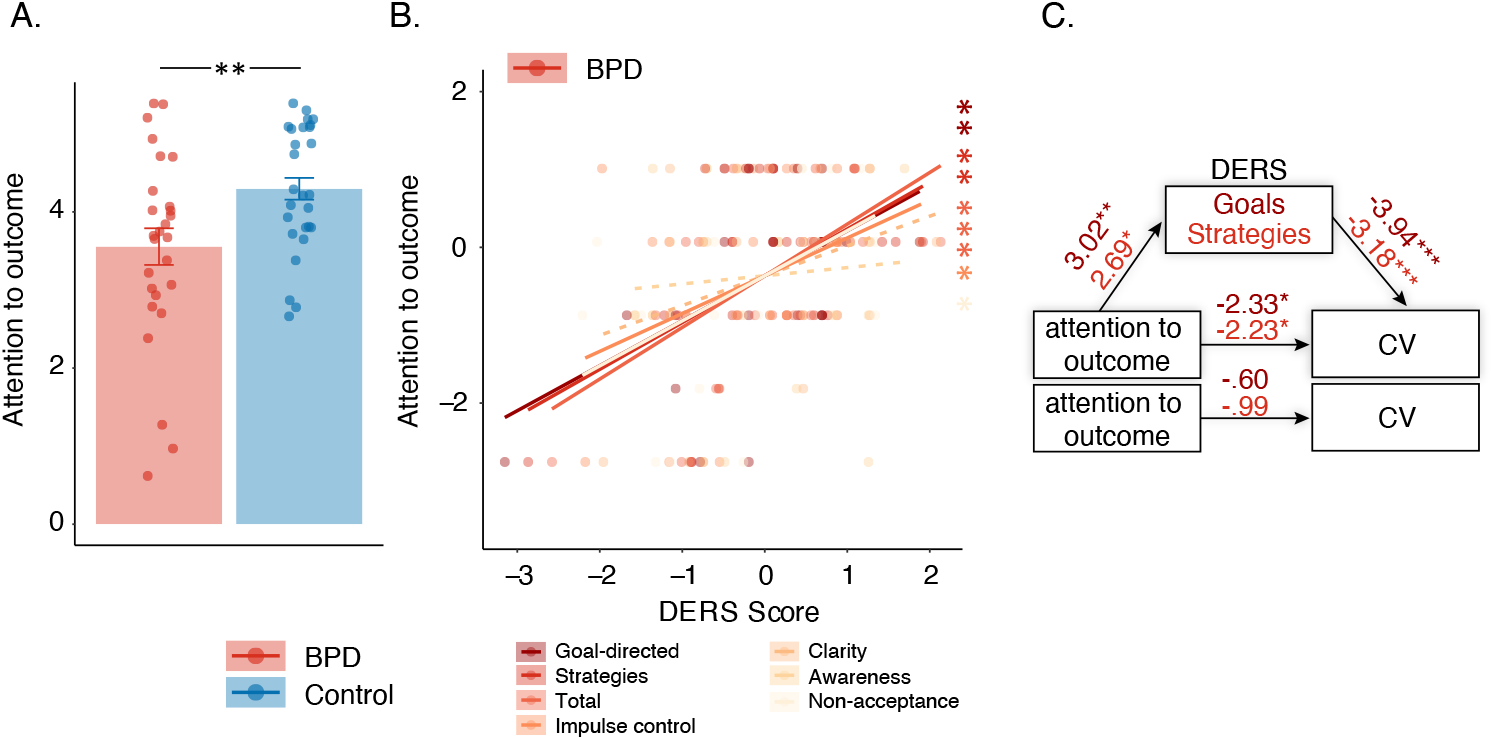
Attention to Outcome During Matching Pennies. (A) Group differences in self-reported attention to outcome. (B) The relationship between attention to outcome and Difficulties in Emotion Regulation (DERS) scores in BPD participants. (C) DERS ‘Goals’ and ‘Strategies’ significantly mediated the relationship between impulsivity and coefficient of variation in saccadic reaction times (CV) in BPD. In A, independent samples T-Test results are reported. In B, z-scores are shown to allow for visualization across different subscales, linear effects results are reported, and solid and dashed lines indicate significant and non-significant regressions, respectively. In C, T-test results are reported, and significance values for indirect effects were obtained using 10000 draws in a bootstrap procedure (mediation package in R; Tingley et al., 2014). In A-B, reported *p* values are Bonferroni corrected. * *p* < .05, ** *p* <.01, *** *p* < .001.

We saw a trend-level main effect of “attention to outcome” on reward rate (Fig. 7; β = .35, *t* = 2.51, *p =* .02, *p_Bonferroni_* = .08), with more attention to outcome associated with marginally increased reward rate. We also found a significant main effect of “attention to outcome” on CV (Fig. 7; β = −.43, *t* = −3.46, *p =* .001, *p_Bonferroni_* = .004), the percentage of anticipatory trials (Fig. 7; β = −.33, *t* = −2.68, *p = .*01, *p_Bonferroni_* = .04) and non-response trials (Fig. 7; β = −.44, *t* = −3.46, *p = .*001, *p_Bonferroni_* = .004), with greater attention to outcome being associated with reduced variation in SRT, fewer anticipatory, and fewer non-response trials (Fig. 7). Finally, we observed a trend level attention to outcome by group interaction on SRT (in absence of main effect; β = .17, *t* = 1.15, *p = .*26, *p_Bonferroni_* = 1.00; interaction β = .77, *t* = 2.40, *p = .*02, *p_Bonferroni_* = .08), and exploratory follow-up tests revealed a significant positive relationship in control participants (Fig. 7B; β = .74, *t* = 2.63, *p = .*01) with more attention to outcome associated with slower SRTs, but a non-significant relationship in BPD (Fig. 7A; β = −.03, *t* = −.20, *p = .*84).

#### Use of a Random Strategy

We observed a significant random strategy by group interaction on the percentage of anticipatory trials (trend level main effect; β = −.25, *t* = −2.10, *p = .*04, *p_Bonferroni_* = .16; interaction β = −.53, *t* = 3.09, *p =* .003, *p_Bonferroni_* = .01), and follow-up tests revealed a significant negative relationship in BPD (Fig. 7A; β = −.52, *t* = −3.41, *p = .*002) with those reporting choosing randomly showing fewer anticipatory trials, and a non-significant relationship in control participants (Fig. 7B; β =.17, *t* = 1.09, *p = .*29). Likewise, we observed a trend-level random strategy by group interaction on CV (non-significant main effect; β = −.21, *t* = −1.63, *p = .*11, *p_Bonferroni_* = .44; interaction β = −.42, *t* = −2.70, *p =* .03, *p_Bonferroni_* = .12), and exploratory follow-up tests revealed a significant negative relationship in BPD (Fig. 7A; β = −.42, *t* = −2.39, *p = .*02) with those reporting choosing randomly showing reduced CV, and a non-significant relationship in control participants (Fig. 7B; β =.13, *t* = .79, *p = .*44).

#### Human vs Computer

We observed a significant main effect of human vs computer on p (wsls) in the color domain (Fig. 7; β = .37, *t* = 2.93, *p = .*01, *p_Bonferroni_* = .04), with those reporting that they would play differently against a human opponent showing increases in WSLS bias. There was no significant group by assessment interaction (β = .002, *t* = .01, *p =* .99, *p_Bonferroni_* = 1.00), and no additional associations with this item and strategic choice behavior (Fig. 7).

#### Correspondence Strategic Assessment, DERS, & BIS

Intriguingly, we found strong positive associations between attention to outcome and the majority of DERS subscales in the BPD group (Fig. 6B & 7A), again with the exception of non-accept and awareness (all *p* < .05, Fig. 6B, 7A). We observed no significant associations between the strategic assessment and the BIS across any subscale (all *p* < .05, Fig. 7).

#### DERS Mediates the Effects of Attention to Outcome on Choice Behavior in BPD (Exploratory Analyses)

We conducted mediation analyses to test whether difficulties in emotional regulation may partially account for the observed relationship between attention to outcome and choice behavior in BPD. We tested for mediation where we saw significant associations between choice behavior and both attention to outcome and DERS scores, and found that the relationship between attention to outcome and CV was significantly mediated by both DERS goals (Fig. 6C; β = −.29, CI [−.60, −.06], *p* = .006) and DERS strategies (Fig. 6C; β = −.22, CI [−.52, −.02], *p* = .03). Relatedly, DERS goals also mediated the trend level relationship between attention to outcome and the percentage of anticipatory trials in BPD (not shown; β = −.22, CI [−.52, −.01], *p* = .03).

## Discussion

In this article, we examined mixed-strategy decision-making in female adolescent outpatients diagnosed with Borderline Personality Disorder (BPD). In particular, we investigated whether adolescents with BPD differed from age and sex matched control participants in their ability to engage in a strategic competition that encouraged randomization in choice patterns (i.e., suppressing choice biases), and whether two core features of BPD, impulsivity and emotional dysregulation, underscored individual differences in choice behavior. We found that the BPD patients as a group showed fewer WSLS biases (Fig. 2), increased variability in reaction times (CV; Fig. 4A) and more anticipatory decisions (Fig. S1A), despite having comparable reward rates (Fig. 3A) relative to control participants. Critically, we found that a subset of BPD participants with high levels of impulsivity and emotional dysregulation showed higher overall reward rates (Fig. 3C & 3D), increased entropy in choice patterns (Fig. 7A), decreased CV (Fig. 4C & 4D), and fewer anticipatory decisions (Fig. S1C & S1D) relative to control (Fig. 3B, 4B & S1B) and BPD participants with lower indices across these domains. In BPD, emotional dysregulation mediated the relationship between impulsivity and CV (Fig. 5). Finally, exploratory analyses revealed that increased vigilance to outcome was associated with increased overall reward rates, decreased CV, and decreased anticipatory decisions (Fig. 7). In BPD, higher levels of emotion dysregulation were associated with increased attention to outcome (Fig. 6B & 7A), and mediated its relationship with CV (Fig. 6C). These results suggest that adolescents with BPD show changes in choice behavior and response variables during Matching Pennies, and that impulsivity and emotional dysregulation contribute to variability in mixed-strategy decision-making in BPD, the latter of which may influence choice behavior, in part, via increasing vigilance to outcome information during the task.

### Group Differences in Matching Pennies Performance

The finding that adolescents with BPD had a greater propensity for variability in reaction times (Fig. 4A) and a greater number of anticipatory trials (Fig. S1A) is in line with prior studies showing increased anticipatory decisions during a stop-signal delay task (Coffey et al., 2011; Nigg et al., 2005), and during the pro- and anti-saccade task (Calancie et al., *in preparation*) in BPD, and could be related to behavioral dis-inhibition (Nigg et al., 2005), although overall response times did not differ between groups (Table 1). Likewise, increased variability (also referred to as ‘increased noise’ in the literature) is a common observance across several psychological illnesses and personality psychopathologies (Kunisato et al., 2012; J. Peters & Büchel, 2011; Robinson & Chase, 2017; Willcutt et al., 2008), including BPD (Kaiser et al., 2008). The increased variation in response time variables in the BPD group could therefore reflect increased variability with emerging psychopathology, and also suggests that our task is sensitive to known behavioral changes in BPD.

We also found *reduced* win-stay, lose-shift (WSLS) bias in BPD relative to controls (Fig. 2), which may have provided a benefit to Matching Pennies performance. In contrast to traditional decision-making paradigms, the strategic nature of our task incentivizes entropy in choice behavior (i.e., fewer reinforcement learning biases) in order to evade exploitation by one’s opponent (Azar & Bar-Eli, 2011; Gauriot et al., 2016; Rapoport & Budescu, 1992; Thaler, 2016). Partial support for this hypothesis comes from the observation that overall reward rate was not significantly lower in the BPD group (Fig. 3A), despite the increases in CV and anticipatory decisions that appeared to considerably serve as a detriment to reward rate and behavior (see previous paragraph and Fig. 7). Reduced WSLS (which was not significantly associated with CV), may have provided a boost to performance by providing less information to the computer opponent to exploit. Although adaptive in this context, this reduced WSLS could be reflective, in part, of changes in limbic processes related to feedback processing (Goyer et al., 1994; Lyoo et al., 1998; Paret et al., 2017; Schuermann et al., 2011; Schulze et al., 2016; Silbersweig et al., 2007; Tebartz Van Elst et al., 2003) and delay discounting in BPD (Barker et al., 2015; Coffey et al., 2011; Ka et al., 2008b; Krause-Utz et al., 2016a; Lawrence et al., 2010; Maraz et al., 2016), as faster discounting of reinforcers may manifest as fewer choice biases. On the other hand, the possibility exists that reduced WSLS could be related to a more deliberative strategy (i.e., suppression of choice biases in an attempt to be random) in the BPD participants. Interestingly, our exploratory analyses revealed a positive association between WSLS and the “human vs computer” item on the strategic questionnaire that assessed whether participants would approach the game differently if playing against a human opponent (Fig. 7). Overall, while BPD patients and controls alike reported that they would indeed play differently against a human (Table 1), those who reported that they would approach the game similarly showed *decreased* WSLS bias, which may have engaged different motivational processes that facilitated the suppression of choice biases and/or randomization over choice options in order to gain strategic advantage.

### Individual Differences in Impulsivity and Matching Pennies Performance

Impulsivity is a tendency to react to external stimuli, often quickly, without fully considering its consequences (Kim & Lee, 2011). We observed differential effects of impulsivity measures in Matching Pennies performance across groups; in the BPD group, higher levels of impulsivity, particularly cognitive instability and motor impulsivity, were associated with *increased* reward rate (Fig. 3C), increased entropy (Fig.7), decreased CV (Fig. 4C) and decreased anticipatory decisions (Fig. S1C). However, these measures were associated with little change among choice variables in control participants (Fig. 3B, 4B & S1B). In particular, two subscales showed this effect: 1) BIS-Cognitive instability, which reflects instability in thought processes (see Results section for more detail; Patton et al., 1995; Stanford et al., 2009); and 2) BIS-Motor, which, despite being classified as impulsive responding in the motor domain, it is worth noting that it is also comprised of several questions relating to impulsive spending behaviors (see Methods section for more detail; Patton et al., 1995; Stanford et al., 2009). Increased impulsivity may diminish the ability to track changing action-outcome contingencies (Kim & Lee, 2011), as has been seen in tasks assessing reinforcement/reversal learning in BPD (Paret et al., 2016, 2017). While this tendency may lead to dysfunction in real-world settings in BPD (Paret et al., 2017), during mixed-strategy games and other related strategic decision-making paradigms, increased behavioral variability (i.e., randomness) can be adaptive (Tervo et al., 2014), and increased impulsivity may therefore have had a facilitatory effect on the ability to produce unpredictable choice sequences.

A few potential (non-mutually exclusive) explanations exist for the discrepancy between associations between impulsivity and choice behavior across groups. First, BIS scores were significantly lower in control participants compared to BPD, and therefore, the lack of associations in the control group could indicate of a floor effect and may suggest that impulsivity levels are too low in this cohort to detect associations with decision-making behavior that might emerge with higher levels of impulsivity. Second, it could suggest that other pathological processes that are associated with BPD are contributing to the observed effect of impulsivity on choice behavior. In support of this notion, we found similar associations among BPD patients with high levels of emotional dysregulation (Fig. 3C, 4C & S1C, see below for discussion), particularly across the goals and strategies subscales, which mediated the association between impulsivity and choice behavior in BPD (Fig. 5), specifically CV. In line with consistent findings in BPD showing exacerbated impulsivity in the face of negative affect and emotional regulation difficulties (Chapman et al., 2008; Koenigsberg et al., 2001; J. R. Peters et al., 2013; Scott et al., 2014; Sebastian et al., 2013), our findings suggests that impulsivity may exert it’s influence on strategic choice behavior in part through the mediating role of emotion dysregulation.

### Individual Differences in Emotion Dysregulation and Matching Pennies Performance

Emotion dysregulation may disrupt self-regulation and deliberative processes in BPD (Hallquist et al., 2018; Sharp et al., 2011), including mentalization (Henco et al., 2020; Sharp et al., 2011). We found that higher levels of emotion dysregulation in the BPD group, particularly, difficulties engaging in goal-directed behavior (goals) and limited access to emotion regulation strategies (strategies), were associated with *increased* reward rate (Fig. 3D), increased entropy (Fig. 7A), decreased CV (Fig. 4D) and decreased anticipatory decisions (Fig. S1D). In particular, two DERS subscales showed this effect: 1) goals, which reflects a lack of ability to engage in goal-directed behaviors when emotionally distressed (see Methods section for more detail; Herr et al., 2013; Gratz & Roemer, 2004) and 2) strategies, which reflects a lack of access to strategies that would allow one to modulate the intensity and/or duration of emotional responses (see Methods section for more detail; Herr et al., 2013; Gratz & Roemer, 2004). In confirmation of the BIS findings in BPD, we also found that DERS impulse control difficulties was marginally associated with higher reward rates (Fig. 4D).

The seemingly facilitatory effect of emotion dysregulation on Matching Pennies performance-related behaviors observed in the current study could reflect modulation of mentalization-related processes in the several potential ways. The first possibility is that given interpersonal challenges in BPD (King-Casas & Chiu, 2012), which may be enhanced with increased emotional dysregulation (Euler et al., 2019), BPD patients with higher levels of emotional dysregulation opted for a more random strategy, and enhanced performance therefore may reflect an increased ability to evade exploitation by the opponent. Studies have shown that knowledge of the task may contribute to variance in behavioral performance in BPD (Paret et al., 2017), thus it is possible that knowledge of the computerized opponent in the current study may have stimulated a change in the strategies adopted by the BPD group; rather than endowing the opponent with human/social qualities that may be more threatening or cognitively laborious, participants with high-levels of emotion dysregulation opted for a more random strategy. In partial support of this hypothesis, we found that higher levels of emotion dysregulation in BPD were associated with marginal *increases* entropy in choice patterns (Fig. 7). Not only would a random strategy potentially result in less exploitation by the opponent, driving increases in reward rate, but could also drive the differences in response variables as individuals may get ‘into a rhythm’ in motor responding using a random strategy (rather than exploitative). Another possibility is that increased emotion dysregulation may have resulted in hypervigilance to the opponent’s actions and outcomes (social information), potentially resulting in enhanced prediction and therefore higher reward rates. In support of this hypothesis, we found that higher levels of emotion dysregulation in BPD were related to increased vigilance to outcome information (Fig. 6B), which inherently carries information about the opponent’s choice (i.e., if the participant as the ‘matcher’ chose the green target and was rewarded, they could infer that the computer chose green that trial as well), which was associated with increased reward rate, decreased CV, and fewer anticipatory decisions, and significantly mediated the relationship between DERS and CV (Fig. 4B). In further support, attention to outcome information was positively associated with the use of a predictive strategy in the BPD group (assessed using our strategic questionnaire; Fig. 7A). Given that we did not observe a negative effect of DERS on strategic choice behavior, our results are inconsistent with the hypothesis that emotion dysregulation resulted in misattribution of the opponent’s actions, but instead suggest that greater difficulties in emotion regulation may contribute to strategic choice behavior by enhancing vigilance to outcome information.

### Conclusions and Next Steps

Economic exchange games are gaining significant traction in understanding deficits in interpersonal functioning across a number of psychiatric conditions (Euler et al., 2019). Game theoretical studies in BPD have found that behaviors during social interactions appear to be *less* modulated by social signals, which in particular contexts, may appear as greater “rationality” in choice (Jeung et al., 2016). While the interpersonal deficits associated with BPD create conflict across a number of domains, in some cases, may have a faciliatory effect on behaving in a self-interested manner in social and economic interactions (Jeung et al., 2016). We found that increased emotional dysregulation and increased impulsivity in BPD may facilitate improvements in strategic choice behavior in this competitive context, although future studies are required to further ascertain whether this is a deliberate process / strategy.

In summary, our results provide a novel account of how two main constructs underlying BPD pathology, impulsivity and emotional dysregulation, affect decision-making in a competitive context. Research supports that individuals with higher clinical indices of impulsivity and affective dysregulation, compared to the BPD clinical group average, experience high therapy-burnout (Yeomans et al., 1994), are more likely to utilize high-cost centers (i.e., ER, inpatient hospitalization) to manage symptoms, and are more likely to attempt suicide (Black et al., 2004; Lieb et al., 2004; Speranza et al., 2011; Yeomans et al., 1994). Despite this, these individuals access the same therapy regimen as other BPD patients (DBT and interpersonal therapy). Perhaps more directed therapeutic interventions that guide decision-making strategies toward the normal curve may correspondingly benefit patient outcomes for this select clinical cohort, similar to trials employing mentalization-based therapies (Bateman & Fonagy, 2010).

In the current study, several aspects of strategic choice behavior were not significantly different across groups, including overall reward rate (Fig. 3A), which could suggest that at the group level, strategic choice behavior remains unaffected in adolescents with BPD. Alternatively, BPD participants, particularly the high impulsivity and emotional dysregulation groups, may have been able to overcome deficits through the use of compensatory strategies (Paret et al., 2017), for example, by increasing vigilance to outcome information (Fig. 6), or by opting for the use of random strategies. A third possibility is that while the current study captured the dynamic and interactive nature of complex, real-world strategic engagements, it only partially captured the *social* components (Parr et al., 2019; Rilling et al., 2002; Vickery et al., 2015), thus may not be sensitive to the full range of interpersonal dysfunction typically associated with BPD. Future studies should therefore seek to understand whether strategic game-play against a human opponent stimulates different mentalization processes in adolescents with BPD. Finally, given that this study was cross-sectional, an important area for future work is to extend this line of inquiry into a large, well-powered, sample using a longitudinal design (potentially in high-risk developmental samples) that explores the relationship between emergent BPD traits in adolescents and functional outcomes through to adulthood.

## Acknowledgements

We thank Ann Lablans, Sean Hickman, and Mike Lewis for outstanding technical assistance, as well as members of the Munoz lab for comments on an earlier draft. We thank Scott Murdison and Jerry Jeyachandra for assistance with aspects of data analysis. This work was supported by a Canadian Institutes of Health Research (CIHR) Operating Grants (MOP-93667 to DPM and FDN-148418 to DPM) and a Southeastern Ontario Academic Medical Organization (SEAMO) AHSC AFP Innovation Fund grant to SKK and DPM. DPM was supported by the Canada Research Chair Program.

## Author Contributions

A.C.P., B.C.C., and D.P.M. designed the experimental protocol; A.C.P. and O.G.C. performed research and collected data; A.C.P., O.G.C. and S.K.K. recruited and screened participants; A.C.P., O.G.C., and B.C.C. analyzed data; A.C.P., O.G.C., D.P.M., and S.K.K. wrote the manuscript.

---

**Figure S1.**Percentage of Anticipatory Decisions during Matching Pennies. (A) Group differences in anticipatory decisions. The relationship between anticipatory rate and BIS impulsivity scores in (B) control and (C) BPD participants. (D) The relationship between anticipatory rate and DERS emotion dysregulation scores in BPD participants. In A, independent samples T-Test results are reported, in B-D, z-scores are shown to allow for visualization across different subscales, linear effects results are reported, and solid and dashed lines indicate significant and non-significant regressions, respectively, and reported *p* values are Bonferroni corrected. * *p* < .05, ** *p* <.01, *** *p* < .001.

