## Supplementary figures and images for "Impulsivity and Emotional Dysregulation Predict Choice Behavior During a Competitive Multiplayer Game in Adolescents with Borderline Personality Disorder"

### Supplemental Figure 1

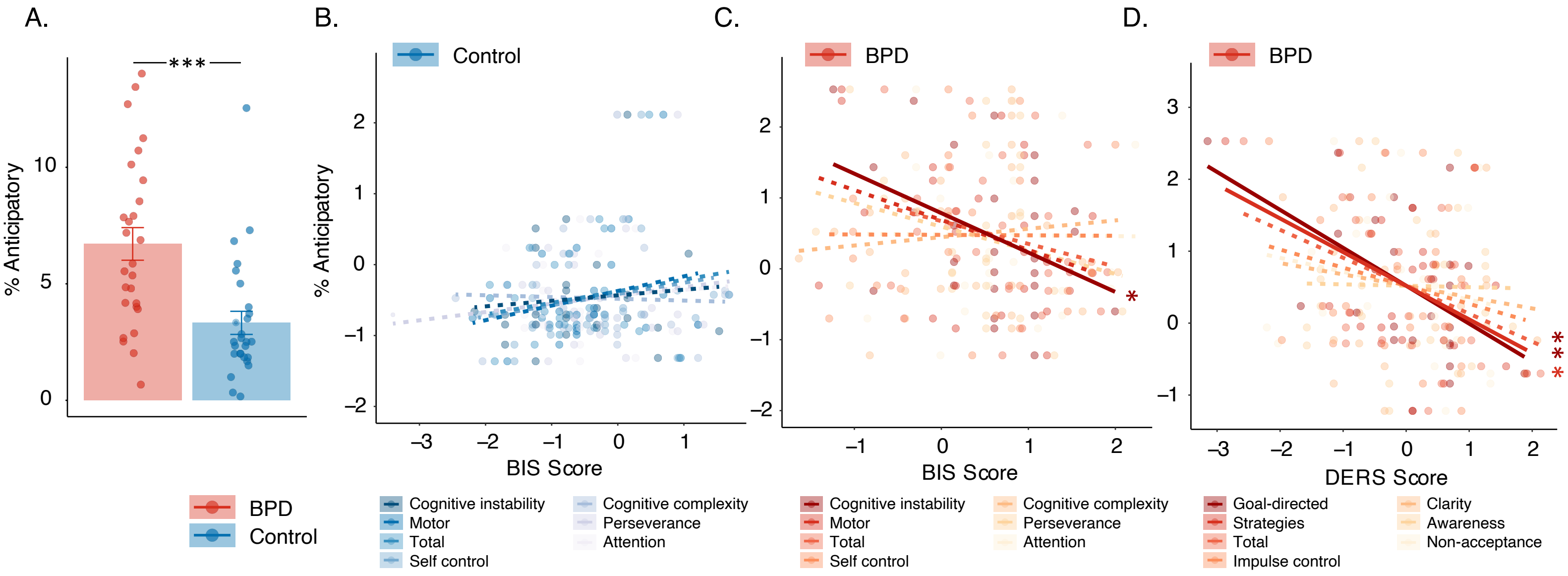


Figure S1. Percentage of anticipatory trials and associations with BIS and DERS scores.
